## Appendix 1 for "Determining which mechanisms underlie facilitation by tussocks in tropical high mountains and their relative importance"

*Appendix: Comparison of the effects of vermiculte and tussocks on soil moisture*

Measuring soil desiccation rates is challenging at high mountains where weather is changing constantly and there are large numbers of visitors. Desiccation rates are usually slow, making long measurement periods necessary. Thus sudden rainfall can ruin two or three days of work. No expensive equipment can be left at the site because of a high risk of vandalism, so researchers need to remain there until a sufficiently-long dry-spell permits obtaining the data. Because of this, we could not measure soils desssication successfully at the study site. Instead, we measured the effects of vermiculite on desiccation rates at a greenhouse. These rates could not be directly compared with those under tussocks or bare soils at our study site. To assess whether desiccation of soil with vermiculite provides a reasonable approximation to what happens under tussocks, in this appendix we will compare the results of our greenhouse experiment with what is expected from theoretical models.

The main driver of desiccation rates is soil temperature (Tovar Romero 2010). We have measurements of the temperature at 2 cm below ground over the day in bare soil and below tussocks using the same equipment described in the main text (Tovar Romero 2010). The measurements were conducted in December, close to the period when most of the plants in vermiculite were still alive. From these data, desiccation rates can be estimated using models from published literature.

Desiccation rates for sandy loams depending on soil temperature at 2 cm below ground where published by Breshears et al. (1998). The soil at the Iztaccíhuatl is also a sandy loam (Galván Díaz 2016). In Breshears et al. (1998), the volumetric soil water content (%), $V$, was accurately predicted by model $V = ae^{-b_T t}$ where $a$ is the water content at the start of the experiment, $b_T$ is the temperature-specific drying rate, and $t$ is drying time in hours[1]. Now, consider that soil water potential $\psi$ changes following the potential function $\psi$ = -c$V^{-d}$ (ICT International 2014). Thus, for a given temperature $T$, $\psi_T = -ca^{-d}e^{b_T dt}$. From this, we can compare the water potential for soils exposed to different temperatures $A$ and $B$ as the ratio $R_{A,B} = \frac{\psi_A}{\psi_B} = \frac{e^{b_A dt}}{e^{b_B dt}} = e^{(b_A - b_B)dt}$. If $A$ is the higher temperature, a value of $R_{A,B}$ = 1.2 would mean that the water potential is 20% lower in the warm soil than in the cool one. This difference in $R_{A,B}$ between both soils increases over time at a rate equal to $D'_{A,B}(\Delta t) = \frac{e^{(b_A - b_B)dt_2}}{e^{(b_A - b_B)dt_i}} = e^{d(b_A - b_B)\Delta t}$ where $\Delta t = t_2 - t_1$, the time interval over which the rate is measured. Because this is a multiplicative rate, we can standardize it per time unit

[1] Note that Breshears et al. incorrectly report that the model is $V = a + e^{-b_T t}$. However, this is at odds with their figure 3a, which shows that $V$ changes linearly with time on a semi-log scale, which is not the case $V = a + e^{-b_T t}$ but holds for $V = ae^{-b_T t}$. Moreover, the latter equation reproduces well the lines and data in figure 3a given the parameters that they report, whereas the incorrect form fails to provide a reasonable fit

as $D_{A,B} = \left[ D'_{A,B} \left( \Delta t \right) \right]^{\frac{1}{\Delta t}} = e^{d\left(b_A - b_B\right)}$. Thus, the rate at which the warmer soil dries relative to a cooler soil is constant over time if temperature remains unchanged. A value of $D_{A,B}$ = 1.1 means that the warm soil dries at a rate 10 % higher than the cooler one.

From our measurements of water potential in drying soils with and without vermiculite we can obtain an empiric estimate of $D_{A,B}$ as

$$D_{A,B} = \left( \frac{\dfrac{\psi_C\left(t_2\right)}{\psi_V\left(t_2\right)}}{\dfrac{\psi_C\left(t_1\right)}{\psi_V\left(t_1\right)}} \right)^{\frac{1}{\Delta t}}$$

where $\psi_C(t)$ and $\psi_V(t)$ are the hydric potentials of soil without (control) and with vermiculite at time $t$. Note that the vermiculite treatment is considered to be analogous to the high temperature treatment in this calculation because vermiculite and low temperatures are expected to slow desiccation. The mean $D_{A,B}$ value estimated from the greenhouse experiment (see figure 2 in the main text) was 1.015 (range 0.994 – 1.085), indicating that control soil is expected to dry at a rate 1.5 % larger than that with vermiculate.

In bare soil, the observed mean maximum temperature was 27 °C, and under tussocks it was 12 °C. Using the estimates for $b_T$ for these temperatures published by BRESHERAS, and a $d$ value of 1.852, which corresponds to sandy soil (ICT International 2014) $D_{A,B}$ value is 1.005, which would suggest that the effect of vermiculite on soil moisture was even larger than the expected effect of tussocks. If we use the $d$ value of 4.228 of clayey soils, $D_{A,B}$ = 1.012, closer to the measured effect of vermiculite, but still slightly smaller. The $D_{A,B}$ for our system would arguably be between 1.005 and 1.012, because the texture of the soil at the study site is intermediate between sandy and clayey.

Our estimates for $D_{A,B}$ at the study site are likely not to be accurate. We are using the maximum temperature in our calculations because the desiccation rates during the nigh are likely to be negligible (Breshears et al. 1998). The use of lower temperature estimates would result in still smaller $D_{A,B}$ estimates. Another factor that we are not considering is the re-wetting of soil during the night due to capillarity and the storage of water in lower soil horizons. This cannot be reproduced in the greenhouse, and would lower our estimates of $D_{A,B}$ from that experiment. It is thus difficult to tell whether vermiculite had the desired effect on soil moisture. However, the available data suggest that it may have provided a reasonable approximation to the behavior of water under tussocks.
